# Hard to catch: Experimental evidence supports evasive mimicry

**DOI:** 10.1101/2020.05.20.102525

**Authors:** Erika Páez V, Janne K. Valkonen, Keith R. Willmott, Pável Matos-Maraví, Marianne Elias, Johanna Mappes

**Author notes:** Contributed equally (Erika Páez V, Janne K. Valkonen).

## Abstract

Most research on aposematism has focused on chemically defended prey, but signalling difficulty of capture remains poorly explored. Similarly to classical Batesian and Müllerian mimicry related to distastefulness, such “evasive aposematism” may also lead to convergence in warning colours, known as evasive mimicry. A prime candidate group for evasive mimicry are *Adelpha* butterflies, which are agile insects and show remarkable colour pattern convergence. We tested the ability of naïve blue tits to learn to avoid and generalise *Adelpha* wing patterns associated with difficulty of capture, and compared their response to that of birds that learned to associate the same wing patterns with distastefulness. Birds learned to avoid all wing patterns tested, but learning was faster with evasive prey compared with distasteful prey. Birds generalised their learned avoidance from evasive models to imperfect mimics if the mimic shared colours with the model. Despite imperfect mimics gaining protection from bird’s generalisation, perfect mimics always had the best fitness, supporting selection for accurate mimicry. Faster avoidance learning and broader generalisation of evasive prey suggest that being hard to catch may deter predators at least as effectively as distastefulness. Our results provide empirical evidence for a potentially widespread alternative scenario, evasive mimicry, for the evolution of similar aposematic colour patterns.

## BACKGROUND

Many organisms with chemical, morphological or behavioural defences often display a conspicuous signal, such as a colour pattern, that warns predators of the potential cost of attacks. Possession of such warning signals is known as aposematism (1,2). In many cases, the effectiveness of aposematism depends on the ability of predators to associate the signal with an unpleasant experience related with the stimulus, and to attribute signal properties to different prey individuals (i.e. generalisation) (reviewed in (3)) (4–6), which results in prey avoidance. Aposematic prey are under positive frequency-dependent selection, which can result in selection for convergence of warning signals among co-occurring defended species, known as Müllerian mimicry (7). Aposematism and Müllerian mimicry associated with distastefulness have been extensively studied in many taxa (8–11), but especially in Lepidoptera (12–16). However, there is increasing evidence that aposematism may also be associated with an alternative defence, namely effective evasiveness (reviewed in (17)). Theoretically, predators should avoid attacking evasive prey since unsuccessful attacks likely represent a significant cost in time and energy to the predator (17–19). In this case, selection exerted by predators is expected to drive convergence in signals that they associate with the evasiveness of their prey (20–25), in a process known as escape mimicry or evasive mimicry (hereafter we use the latter term).

Previous experiments have shown that bird predators can use visual cues to identify evasive prey (26–28), but more empirical work is needed to test whether outstanding potential examples of evasive mimicry could indeed be the result of selection for such signals related to evasiveness. One such example is the diverse Neotropical butterfly genus *Adelpha*, where repeated convergence of their apparently conspicuous and contrasting wing patterns among distantly related sympatric species has been interpreted as evidence for mimicry (29–31). Mimicry in *Adelpha* has been hypothesized to be at least partly driven by chemical defences in some species (32–34), but there is currently limited, conflicting evidence for distastefulness (22,33,35,36). In contrast to most classic groups of chemically defended butterflies, *Adelpha* butterflies have short and stout thoraxes and exhibit fast and unpredictable flight (K.W., personal observations) (19), which are favourable traits for butterflies aiming to escape predators (35,37), making the genus a prime candidate for evasive mimicry (38).

In this study, we used models of wing patterns of *Adelpha* butterflies and wild blue tits as naïve bird predators to address the following questions: 1. Can birds learn to associate naturally occurring wing patterns with evasiveness of prey? 2. Can such a signal be generalised across putative mimetic species? 3. What type of secondary defence drives faster learning by predators, evasiveness or distastefulness?

## MATERIALS AND METHODS

We used wild blue tits (*Cyanistes caeruleus*) to examine whether birds can learn to avoid *Adelpha* colour patterns associated with evasive (escaping) behaviour, and whether birds generalised the learned avoidance across similar, naturally occurring wing patterns. In addition, we conducted parallel experiments with distasteful prey having the same colour pattern but not evasiveness.

Experiments were conducted from January to March 2019 at Konnevesi Research Station in Central Finland, which provided the infrastructure, wildlife research and collection permits, and expertise needed to conduct experiments with wild birds in captivity. Blue tits were captured from feeding sites around the station and were maintained in captivity for a maximum of 10 days, during which time they were kept singly in illuminated plywood cages (daily light period of 12 h 30 min) with food and fresh water available *ad libitum*. After experiments birds were ringed and released into the site of capture.

### Artificial prey

Artificial defended prey (4.1 × 2.5 cm) were constructed by printing images (HP Color Laserjet CP2025, regular printer paper) of different wing colour patterns displayed by the species *Adelpha salmoneus*, *A. cocala*, and *A. epione* (figure 1). These species represent three putatively distinct mimicry rings (29,31) and were chosen because they differ in colour and pattern, to enable us to test if apparently distinct signals may provide protection from predation in evasive mimicry. An entirely dark brown model of a non-defended prey was constructed as a control. To make prey attractive for birds, a piece of almond (reward), was glued to the underside of prey. For distasteful models, almonds were soaked in chloroquine phosphate solution (7%), to give them a bitter taste (following e.g., (39)).

**Figure 1.**
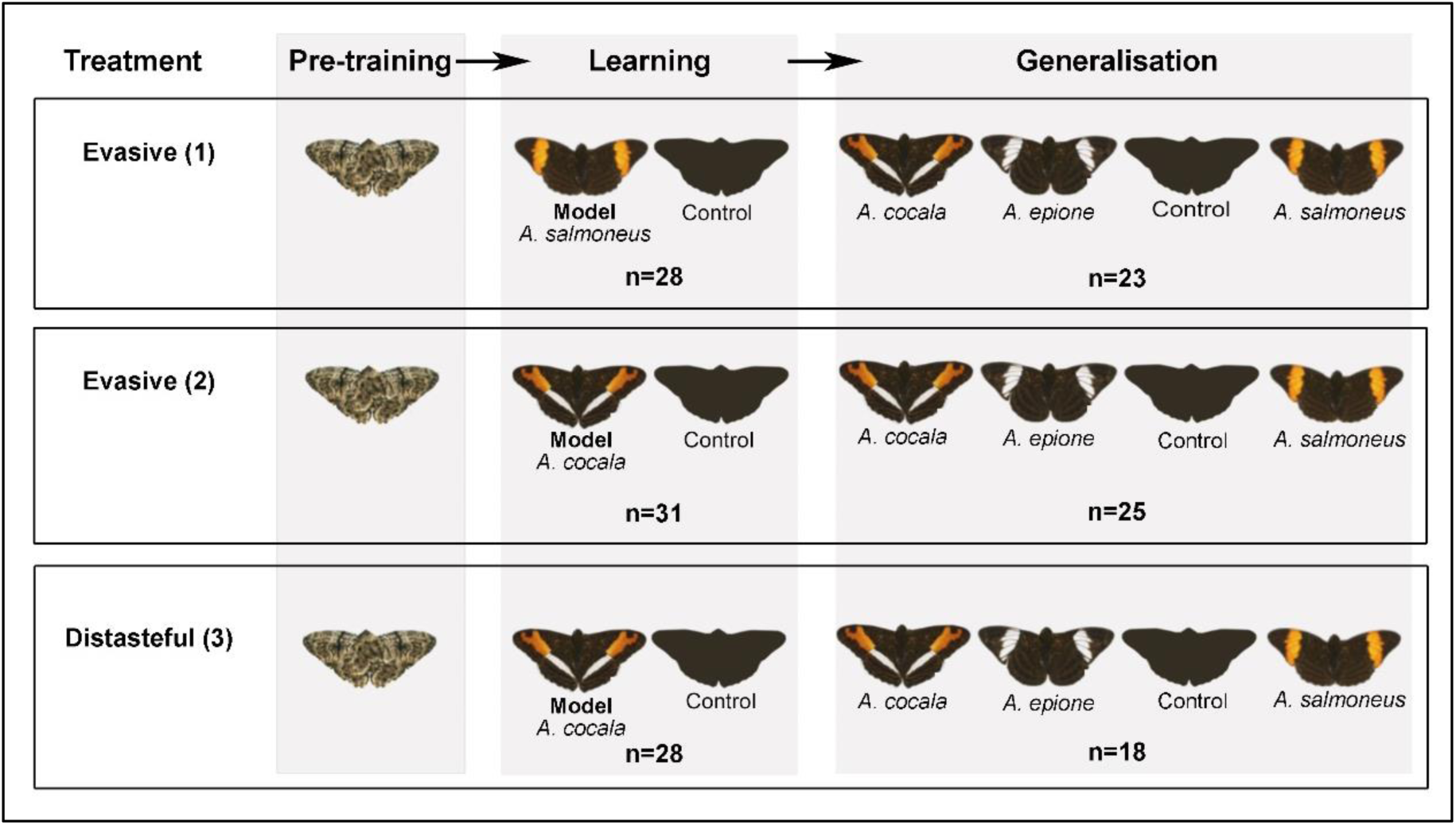
Schematic illustration of the experimental design that consisted of 3 phases: pre-training, learning and generalisation. A forewing orange-banded prey (*A. salmoneus)* was presented as a model for group 1, and as an imperfect mimic during generalisation for group 2 and 3. A transverse forewing orange/hindwing white-banded prey (*A. cocala*) was the model for group 2 and 3, and an imperfect mimic during generalisation for group 1. The forewing white-banded prey (*A. epione*) was presented as an imperfect mimic during generalisation for all groups.

### Experimental procedures

The experiment took place in experimental aviaries of 49 × 48 × 67 cm illuminated by light bulbs. Each aviary contained a perch and a water bowl. Birds were observed through a one-way glass situated on the front of the aviary. Two plastic prey holders gliding on aluminium profile rails (fixed on both sides of the aviary’s floor) allowed simulation of the artificial prey’s escaping (electronic supplementary material, S1 figure 2).

**Figure 2.**
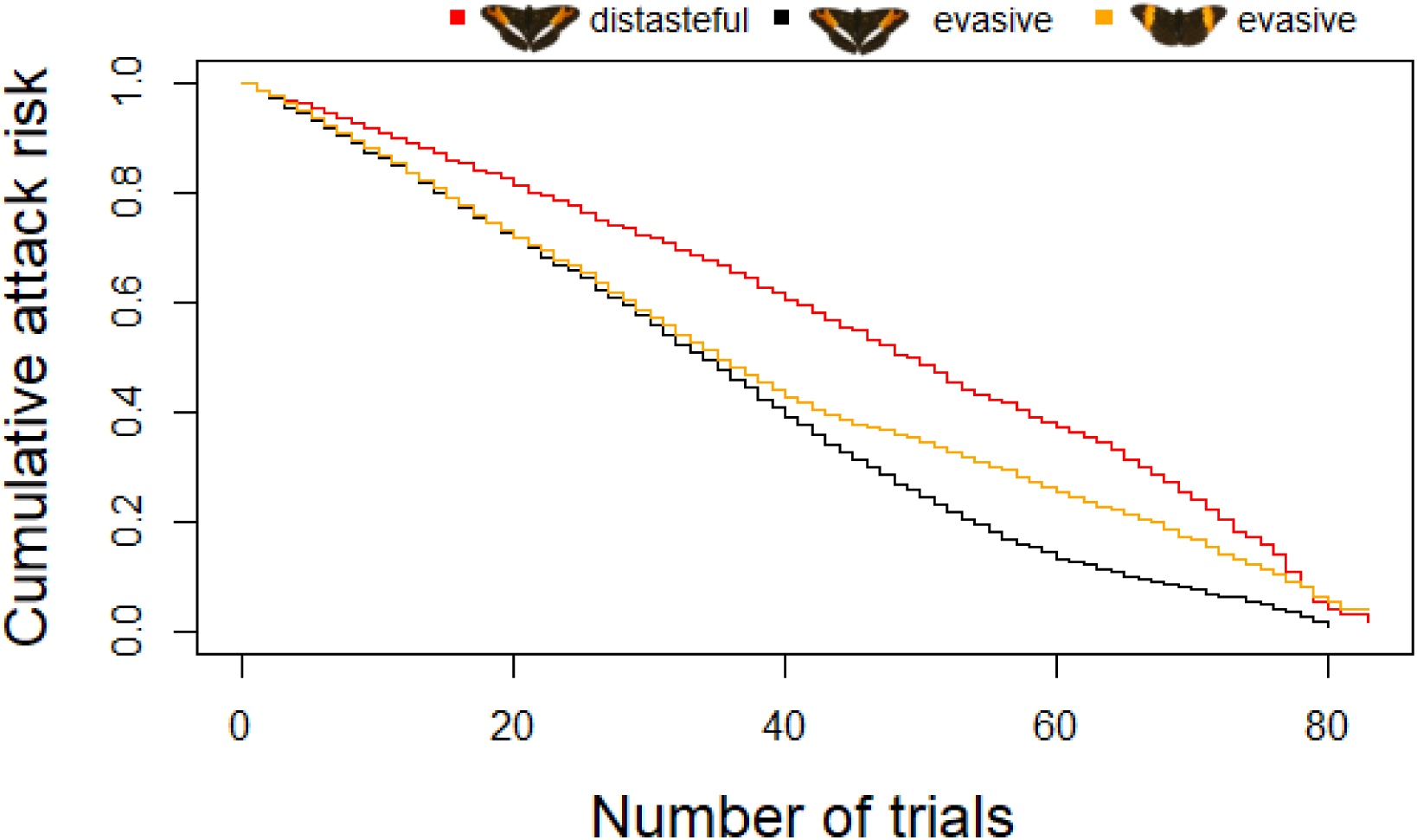
Cumulative attack risk curves of the model during learning trials. Each line represents a type of prey model presented to each group of birds (group 1 yellow line, group 2 black line, and group 3 red line.

#### Avoidance learning

We used 87 birds, trained to attack on artificial butterflies (see the electronic supplementary material, S1 for details of the training procedure), and divided into 3 treatment groups (figure 1). The first two groups were trained to avoid evasive prey and a third group was trained to avoid distasteful prey with the same colour wing band pattern as group 2. Before initiating the experiment, birds were habituated to the experimental aviary for at least an hour. Experiments consisted of a series of trials where two prey were presented simultaneously to the bird. Birds learning evasive model prey had one opportunity of attack per trial, and they were allowed to capture and eat only the control prey, whereas the evasive prey was always rapidly pulled out of reach (i.e., escaping) when attacked. In the treatment group where birds were trained to avoid distasteful prey they were allowed to consume the attacked prey (i.e., distasteful prey and control prey). Training presentations continued for at maximum 80 trials or until the bird attacked an evasive or distasteful prey no more than twice over ten consecutive trials. This learning criterion ensured that all birds reached the same level of learning, which was important for the following generalisation test.

#### Generalisation of learned avoidance to other prey (imperfect mimics)

We used only birds that achieved the learning criteria in previous phase (group 1 n=23, group 2 n=25 [number of birds that learned is 29 out of 31, but data of four birds that followed a different preliminary protocol for generalisation are not included], group 3 n=18) to test whether and to what extent the previously learned avoidance of warning coloration associated with evasiveness (group 1 and 2) or distastefulness (group 3) was generalised to novel wing colour patterns that shared some features with the learned colour pattern (i.e., either colour or pattern). Those novel colour patterns are referred to as imperfect mimics. Birds were simultaneously presented with four types of prey: a (i) control prey, (ii) the model they had learned and (iii) two imperfect mimics (figure 1; see electronic supplementary material, S1 for detailed description).

### Statistical analyses

#### Avoidance learning

We examined whether birds from group 1 (n=28) and group 2 (n=31) learned to avoid wing patterns associated with evasiveness, and whether wing colour pattern affected learning speed. We used a mixed-effects Cox regression model (“coxme” package version 2.2.10 in RStudio v.3.5.3; RStudio 2019) where the response variable was the survival probability of the control prey within trials. The wing colour pattern that the bird learned to avoid as the evasive model was added as an explanatory factor, and bird individuals as a random effect.

#### Comparison of avoidance learning between evasive and distasteful prey

To compare avoidance learning among birds facing aposematic prey signalling for evasiveness, and birds facing aposematic prey signalling for distastefulness with the same colour pattern (group 2 and 3, respectively; figure 1), we performed another mixed-effect Cox regression model. The response variable was the survival probability of the control prey within trials and the explanatory variable was the type of prey defence (i.e. evasiveness or distastefulness). Bird individual was defined as a random effect.

#### Generalisation of learned avoidance to other prey (imperfect mimics)

For each experimental group, to test for differences in attack probabilities between the different types of prey (the control, the model and the two imperfect mimics, figure 1), we calculated the log-likelihood of observing the number of attacks that were recorded on each prey type across all in the group as follows.

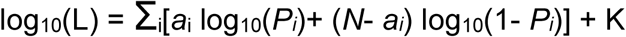

Where *i* is one of the four prey types; *N* is the total number of trials; *a_i_* is the number of times a butterfly of type *i* was attacked; *P_i_* is the attack rate of butterflies of type *i* and K is a constant term that disappears in model comparisons.

We explored several scenarios where attack rates of different types of prey could be equal or not (see electronic supplementary material, S3 for a list of all those scenarios), and calculated the log-likelihood functions of those scenarios. As an example, a scenario where the attack rate on the control is equal to those on the imperfect mimics and higher than that on the model means that birds do not generalise the learned avoidance to the imperfect mimics; a scenario where the attack rate on the model is equal to those on the imperfect mimics and lower than that on the control means that birds have fully generalised the learned avoidance to the imperfect mimics; and a scenario where the attack rate on the imperfect mimics is lower than that on the control but higher than that on the model means that birds have partially generalised the learned avoidance to the imperfect mimics.

Models were selected on the basis of their AICc, which accounts for the number of parameters and the sample size. For each group, the model with the lowest AICc was considered the best. We considered that models within a 2-unit AICc interval from the best model could not be rejected.

## RESULTS

### Avoidance learning

According to the learning criterion, most birds learned to avoid their evasive prey model: 23 out of 28 birds from group 1 (i.e., orange forewing band) and 29 out of 31 birds from group 2 (i.e., transverse orange/white band). Additionally, 18 out of 28 birds (group 3) learned to avoid the distasteful prey model.

The mixed-effects Cox regression model detected no significant differences (*Z=*0.05; *P*=0.96) between birds that learned to avoid different wing patterns of evasive prey (i.e. group 1 and group 2), but showed that birds learned to avoid evasive prey (group 2) significantly faster than distasteful prey with the same wing pattern (group 3) (*Z*=-3.21; *P*=0.001) (figure 2).

### Generalisation of learned avoidance to other prey (imperfect mimics)

Bird’s attack frequencies on mimics differed within and among groups (figure 3; electronic supplementary material S2). For group 1 (prey with orange forewing band as evasive model), in the best scenario learned avoidance was fully generalised to the imperfect mimic that shared a colour (orange) with the model, while the other imperfect mimic (white forewing band) was attacked as much as the control (estimated attack rates on the model and the orange/white mimic: 0.109; estimated attack rates on the control and the white mimic: 0.391; AICc = 45.079, electronic supplementary material S3). Two additional scenarios could be considered as similarly plausible, based on their AICc. One was similar to the previous, except that the orange/white mimic was attacked more often than the model (but still less than the control; estimated attack rate on the model: 0.043; on the orange/white mimic: 0.174; on the control and white mimic: 0.391; AICc = 46.809, electronic supplementary material S3), indicating partial generalisation.

**Figure 3.**
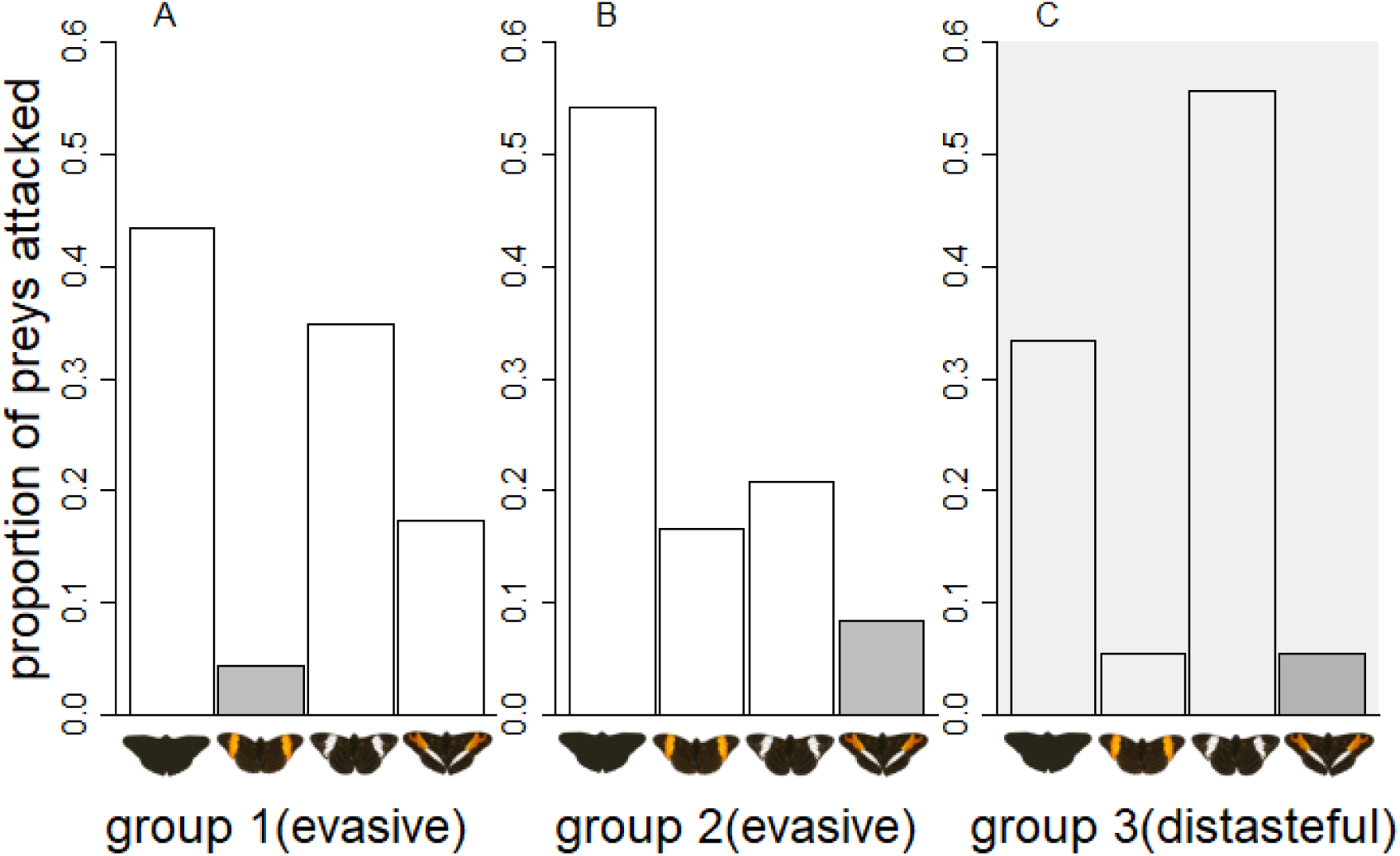
Comparison among observed attack rates on the model, control and imperfect mimics during generalisation tests. Bars illustrate the attack proportion within groups on different wing colour patterns after birds learned to avoid the model pattern (group 1: n=23, group 2: n=25, group 3: n=18). The grey bar indicates the model.

In the other, only the model was attacked less than the control, implying no generalisation (attack rate on the model: 0.043, attack rates on the control and mimics: 0.319; AICc = 45.690, electronic supplementary material S3).

Regarding group 2, (orange/white as evasive model), in the best scenario avoidance was fully generalised to both imperfect mimics, which both shared a colour with the model (estimated attack rate on the model and mimics: 0.188, estimated attack rate on the control, 0.435, AICc = 47.732, electronic supplementary material S3). Two additional scenarios were within a 2-unit AICc interval with that of the model. One of those scenarios was similar to the previous, except that generalisation to the mimics was partial (estimated attack rate on the model: 0.080, estimated attack rate on the mimics: 0.261, estimated attack rate on the control: 0.560, AICc = 49.481, electronic supplementary material S3). In the other, generalisation only applied to the orange mimic, which suffered an attack rate similar to that of the model (estimated attack rates on the model and orange mimic: 0.109; estimated attack rates on the control and white mimic: 0.391, AICc = 49.415, electronic supplementary material S3).

In group 3 (orange/white as distasteful model), a single scenario stood out as best, in which avoidance was fully generalised to the orange mimic only (estimated attack rates on the model and orange mimic: 0.109; estimated attack rates on the control and white mimic: 0.391, AICc = 33.517, electronic supplementary material S3).

## DISCUSSION

### Learning and generalisation of signals associated with an effective escaping ability

The idea that some butterflies have evolved signalling of evasiveness as an anti-predator defence has a long history (19,21,40–42). Still, surprisingly few experiments to date have tested the idea (22,26–28), and nobody has so far tested whether predators can remember and generalise their learned avoidance to other species with signals that are similar to some extent, which is crucial for the evolution of mimicry. Gibson (26,27) and Hancox & Allen (28) presented wild avian predators with artificial prey (i.e. dyed millet seeds, coloured mealworms or pastry models) that disappeared from sight when attacked. After extensive training (approx. 20 days), it was observed that birds reduced their attacks on such hard-to-catch prey. We showed that wild birds, with no experience of *Adelpha* butterflies, were able to associate both orange and orange/white patterns with evasiveness within a day of training. Unlike previous experiments (26–28), our birds faced a “simpler” prey scenario (39), with a warningly coloured prey that could be easily discriminated from the non-defended prey, which may explain the faster avoidance learning we observed.

Our results also showed that birds were able to generalise their learned avoidance to novel, somewhat similar prey (i.e., that shared either a colour or the pattern with the learned model), even though perfect mimics were always the most strongly avoided. Previous work on distasteful prey found that avian predators primarily focus on colour, rather than pattern (43–47) or wing shape (48), when learning and generalising aposematic visual signals. Our findings seem to be consistent with these studies, because all three groups of birds generalised their avoidance to evasive prey that presented a colour in common with the formerly learned model, despite harbouring different patterns, and in group 1, where one imperfect mimic shared the pattern but no colour with the model, birds did not generalise to that mimic. Moreover, although we did not formally test whether some colours or patterns are more efficient as a visual cue for learning or generalisation, we sometimes observed an asymmetrical generalisation (e.g., higher attack rate on the white than the orange mimic in group 3). Further experiments comparing models with different colours could shed light on whether some colours are better learned than others.

The three *Adelpha* species we studied are not regarded as strongly co-mimetic, since a number of other species show much more similar (practically identical) colour patterns, concordant geographic variation and broader sympatry (29). Preliminary trials in our experiment suggested that our predator was incapable of distinguishing among the most perfect co-mimics of *A. cocala*, so we expanded our experiment to include more dissimilar species to examine the significance of mimetic accuracy. Although imperfect mimics gained varying levels of protection, the perfect mimic appeared to mostly be the best protected. Those results suggest strong selection on mimetic fidelity and could explain the extremely close similarity within some putative *Adelpha* mimicry rings.

### Evasiveness versus distastefulness as deterrents to predators

Learning about distastefulness is thought to be generally quicker and easier than evasiveness because prey unprofitability can be determined, unambiguously, from a single experience when prey is ingested. By contrast, a prey individual might escape capture because of better escaping ability, or just because of chance (17). There is thus some disagreement about the circumstances under which evasive aposematism and mimicry might occur and the extent to which its evolution might be different from that of aposematism and mimicry based on distastefulness (6).

In our experiments, in contrast to expectations (17), birds learned to avoid evasive prey faster than distasteful prey, and learning seemed to be easier as a higher proportion of birds achieved the learning criterion with evasive prey (94%) compared to distasteful prey (63%). The close spatio-temporal association between the unrewarding experience (loss of prey) and the received signal could help predators to learn faster about evasiveness, which might not always be the case for distasteful prey (e.g., delayed emetic effect when ingesting a toxic prey, (49)). There is also the possibility of significant variation in distastefulness even within the population of a single mimicry ring as a result of differences in larval host plants and access to adult resources (50,51), or intra and interspecific variation in a predator’s tolerance to distastefulness (49,52–55). Therefore, signals associated with prey evasiveness may actually provide a more reliable message to birds about unprofitability than does aposematic signalling related to toxicity.

Another potential explanation for faster learning is that the decision to attack presumably reflects a trade-off between costs and benefits. All toxic prey also contain nutrients (56) and many birds handle such prey by removing the most toxic body parts (57) or simply make a strategic decision to eat a certain amount of toxin to simultaneously acquire nutrients (58). When a bird predator is hungry, the cost of eating something distasteful might be lower than the benefits of achieving their nutritional needs from a defended prey (59), and the cost of pursuing a prey that is impossible to catch will be increased. In other words, hungry birds may prefer to pursue a somewhat distasteful prey providing at least limited nutrients rather than an evasive prey providing no nutrients.

We also observed dissimilar generalisation patterns between evasive and distasteful treatments, suggesting wider generalisation among colour morphs when the prey defence is evasiveness. Groups 2 (evasive treatment) and 3 (distasteful treatment) had the same model (orange/white transverse band). In group 2 (evasive treatment), in two out of three best scenarios birds generalised to some extent their learned avoidance toward the prey sharing any wing colour with the model, and both imperfect mimics were attacked less than the control. By contrast, in group 3 (distasteful treatment), birds only avoided the orange imperfect mimic, as the white imperfect mimic was highly attacked, despite the fact that the white colour was also present in the model. It has been suggested that selection for accurate mimicry can be affected by different factors (6) such as level of prey distastefulness or unpleasantness (56,57). Broad generalisation to imperfect mimics was observed in previous studies when the model was highly distasteful or unpleasant (see in (60)). Our results, along with those showing faster avoidance learning with evasive prey, suggest that evasiveness is another powerful dimension of defence that affects a predator’s decision whether to attack warningly coloured prey. More experiments with different types of predators and signals are nevertheless needed to examine whether generalisation tends to be broader across mimics where the model is defended by evasiveness rather than distastefulness or toxicity.

## CONCLUSION

Although distastefulness has been considered as a prime adaptive defence mechanism against predation in aposematic butterflies, evasiveness is also likely to be important in a number of other groups. Our results give a strong experimental support for the hypothesis, previously mostly based on field observations, that predators can learn and generalise naturally occurring colour pattern signals that are associated with the escaping ability of prey. We therefore argue that evasive mimicry is a plausible explanation for colour pattern convergence in fast moving prey, such as the *Adelpha* butterflies that are the subject of this study.

### Ethics

The Southwest Finland Centre for Economic Development, Transport and Environment (VARELY/294/2015) and National Animal Experimental Board (ESAVI/9114/04.10.07/2014) provided permission to capture and keep wild blue tits (*Cyanistes caeruleus*) in captivity and to use them in behavioural studies.

## Data accessibility

## Author’s contribution

JM, KRW, ME and PMM conceived the project. JM, EPV, JV, designed the experimental setup, with input from KRW and ME. EPV, JV, PMM and JM ran the experiments. EPV, JV and ME performed statistical analyses. All authors discussed the protocol and results throughout the study. EPV wrote the paper with contributions from all authors. All authors gave final approval for publication and agree to be held accountable for the work performed therein.

## Acknowledgments

We acknowledge Helinä Nisu for taking loving care of birds and Konnevesi Research Station for facilities to conduct this experiment. EPV and ME thank Christine Calvas from the Center of Ecology and Conservation Sciences for kindly handling the funding that allowed EPV to travel to Konnevesi. EPV thanks to Sebastián Mena for designing figure 2 (electronic supplementary materials). PMM thank the support of the Department of Ecology (Biology Centre, CAS) and its head, Vojtěch Novotný, for partly covering travel expenses to carry out the behavioural experiments. We thank the organizers of the 8th International Conference on the Biology of Butterflies, held in Bangalore, India, in 2018, where the idea of this project emerged.

## Footnotes

Electronic supplementary material is available online at

