## Supplementary material for "Hard to catch: Experimental evidence supports evasive mimicry": S1

### S1. Detailed protocol description

#### *Pre-training phase*

The day before experiments, birds were trained in a stepwise manner to handle an artificial prey item bearing a cryptic colour pattern (figure 1e). Pre-training took place in the birds' home cages and started when lights were turned on at 8:40 am. Food trays were removed from the home cages and birds were allowed to eat four sunflower seeds to start the pre-training, which lasted the whole day.

Birds had to accomplish a sequence of 4 tasks to finish this phase. First, they had to consume 4 pieces of almond, placed above pinned pre-training prey. Second, birds had to consume the almond that was under the pre-training prey but half visible. Third, birds had to find the almond that was not visible anymore unless prey was flipped upside down. Finally, the almond was completely hidden (a glued square of paper covered it), and so birds had to rip the paper in order to find the reward.

#### *Experimental procedures*

We used 90 birds, divided into 3 treatment groups depending on the prey wing colour pattern and which unprofitable feature (i. e., evasiveness or distastefulness) was being taught to be avoided. Group 1: evasive prey bearing an orange forewing band wing pattern (n=30) (figure 1a). Group 2: evasive prey bearing a transverse orange forewing and white hindwing band wing pattern (n=31) (figure 1b). Group 3: distasteful prey bearing the same wing pattern as in group 2 (n=29) (figure 1b). For the generalisation phase (see below) we used the white forewing band wing pattern (figure 1c) for all groups.

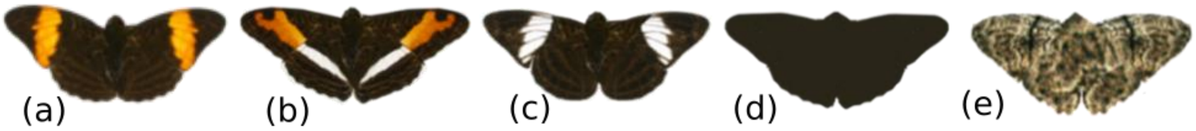


*Figure 1. Artificial prey items*

*Avoidance learning of evasive prey*

Birds were tested individually in the experimental aviary. The day of the experiment, birds were habituated to the experimental aviary (see figure 2) for at least an hour during which they had to eat in a stepwise manner: two sunflower seeds (each one situated next to the aluminium rails), and then two pre-training prey. If birds ate both sunflower seeds and pre-training prey, it was considered they were ready to start the experiment.

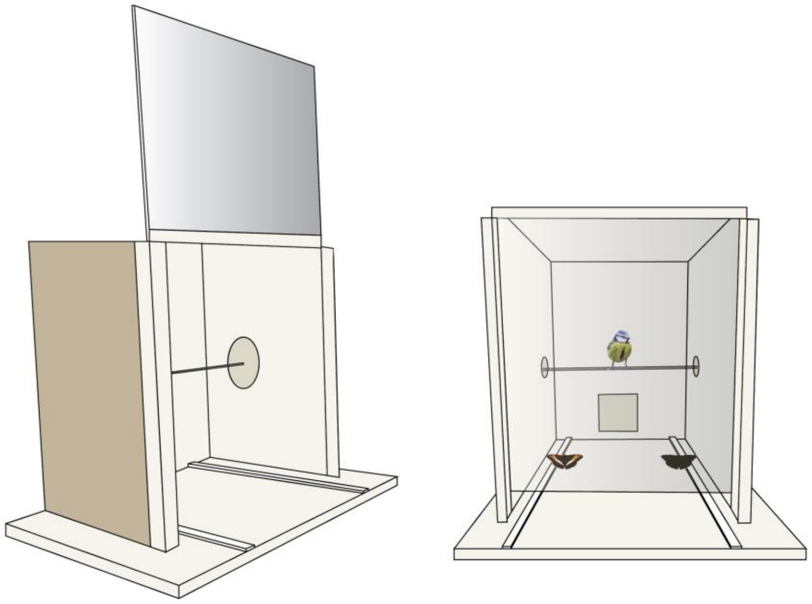

*Figure 2. Experimental aviary*

Each trial consisted of simultaneously presenting to the bird (and in alternating positions per trial), one *non-evasive* prey with the control colour pattern and one *evasive* prey with an *Adelpha* wing colour pattern according to the treatment group. Each trial time was set to a maximum of 3 minutes.

To make sure that each bird had the opportunity of experiencing both types of prey (i.e. *evasive* and *non-evasive* prey), no prey was removed until attacked during the first five trials. After this, the bird had only one opportunity of attack per trial. The bird was allowed to capture and eat only the *non-evasive* prey (i.e., control), whereas the *evasive* prey was always rapidly pulled out when attacked. If the bird did not attempt to capture any of the prey for 3 minutes from when the bird first saw both prey, it was offered a *pre-training* prey to monitor the bird's motivation to continue foraging. If the bird did not attack the *pre-training* prey, a sunflower seed was offered to avoid starvation and then it received a 10-minute break without any food. After the break, another *pre-training* prey was offered and if the bird attacked it, presentations of *evasive* prey and *non-evasive* prey continued until it learned avoidance (see below), or when the bird reached a maximum of 80 presentations. Based on preliminary trials designed to optimize the experiments, we considered the bird to have learned to avoid the *evasive* prey when it did not attack this prey (and thus attacked the *non-evasive* prey) more than twice over ten consecutive trials. Following this learning criterion, "quick" learners (i.e., those that learn after a small number of trials) encountered fewer prey than "slow" learners, although all birds had the same level of learning, which was important for the generalisation test (see below).

When birds had completed this experiment, they had a break of at least 2 hours with 3 sunflower seeds offered each 30 minutes before starting the generalisation test (see below). Birds that took longer to learn thus finished experiments late in the afternoon (around 17h00) and were placed back in their home cages with food *ad libitum* and water until the next day to continue with the generalisation experiment.

*Avoidance learning of distasteful prey*

We conducted in parallel both learning and generalisation tests (see below) similarly to evasive prey treatments (group 1 and 2), except that distasteful prey were substituted for evasive prey. Only the transverse forewing orange/hindwing white band pattern was tested.

*Generalisation of evasive or distasteful prey*

If birds achieved the learning criteria, a 45-min to 1 hour break was set until generalisation phase get started. Only for birds that end the learning phase after 17h00, generalisation phase started early in the following morning. For this test, four types of prey were presented to birds in a T-shaped tray (figure 3). On average, each bird received a 15-minute habituation period to the new type of tray, during which three *pre-training* prey with one sunflower seed were offered. When all *pre-training* prey and the sunflower seeds were consumed, it was considered that the bird was ready to start the test. We simultaneously presented all prey: (i) control, (ii) the model (i.e., wing colour patterns that birds were trained upon during learning phase), (iii) two imperfect mimics novel to the birds which could show the same colours but not pattern (i.e., transverse forewing orange/hindwing white band for group 1; forewing orange band and forewing white band for group 2 and group 3), and only for group 1 the imperfect mimic that had the same pattern but different colour (i.e., forewing white band) than the model was also tested. The first choice of attack was registered for each bird.

A preliminary test using a species that closely resembled the model showed that birds were incapable of distinguishing between these two species, so for the generalisation experiments we used the same pattern during the learning phase and we introduced patterns that differed either by colour or pattern from the model.

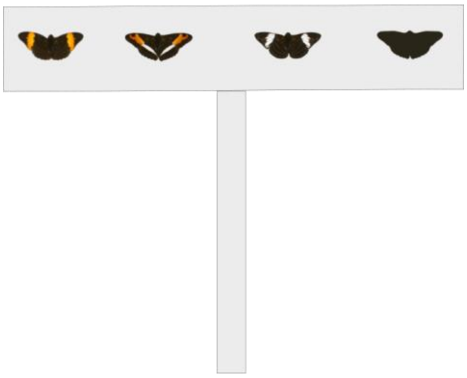

*Figure 3. Illustration of prey presentation during the generalisation test*
