## Supplementary material for "Hard to catch: Experimental evidence supports evasive mimicry": S2

**S2. Frequency of prey attacked in the generalisation test for each experimental group** Numbers in bold correspond to the attack counts on each pattern that was used as a model in the learning experiment. Birds from group 1 (evasiveness) learned to avoid the orange band pattern; birds from group 2 (evasiveness) and 3 (distastefulness) learned to avoid the orange/white band pattern.

| | Attacks on wing patterns presented during generalisation test | | | |
|---|---|---|---|---|
| **Wing patterns learned to avoid** | 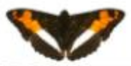 | 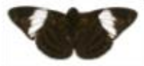 | 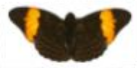 | 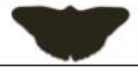 |
| group 1 - evasive prey 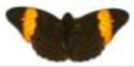 | 4 | 8 | **1** | 10 |
| group 2 - evasive prey 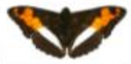 | **2** | 5 | 4 | 14 |
| group 3 - distasteful prey 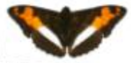 | **1** | 10 | 1 | 6 |
