## Supplementary material for "Hard to catch: Experimental evidence supports evasive mimicry": S3

**S3. Generalisation tests: scenarios investigated and results**

The scenarios investigated are:

(1) All attack rates are equal: $P_c = P_{ow} = P_o = P_w$. This scenario indicates no memorability of avoidance and no generalisation.

(2) All attack rates are different: $P_c \neq P_{ow} \neq P_o \neq P_w$. This scenario indicates partial and unequal generalisation on the two imperfect mimics, if $P_c$ is highest and $P_o$ for group 1 or $P_{ow}$ for groups 2 and 3 is lowest.

(3) Only the attack rate on the control is different: $P_c \neq P_{ow} = P_o = P_w$. This scenario indicates full generalisation to both imperfect mimics if $P_c < P_i$ for each $i \neq$ c.

(4) Only the attack rate on the model is different: $P_o \neq P_c = P_{ow} = P_w$ for group 1 and $P_{ow} \neq P_c = P_o = P_w$ for groups 2 and 3. This scenario indicates no generalisation to imperfect mimics if the attack rate of the model is higher than that of the other types.

(5) The attack of the control and of the white imperfect mimics are equal, and the attack rate of the model and the other imperfect mimic are equal, and different from that of the control: $P_c = P_w \neq P_o = P_{ow}$. This scenario indicates full generalisation to the imperfect mimic that shares the orange colour with the model but not to white imperfect mimic if $P_c > P_o$.

(6) The attack rates of the control and the model are different; the attack rates of the two imperfect mimics are equal and different from those of the control and the model: $P_c \neq P_o \neq P_{ow} = P_w$ for group 1 and $P_c \neq P_{ow} \neq P_o = P_w$ for groups 2 and 3. This scenario indicates partial generalisation to both imperfect mimics, if the attack rate of the control is highest and that of the model is lowest.

(7) The attack rate of the control and the white imperfect mimic are equal, and the attack rates of the orange and orange/white are different, and different from that of the control: $P_c = P_w \neq P_o \neq P_{ow}$. This scenario indicates partial generalisation to the imperfect mimic sharing the orange colour with the model, if the attack rate on the control is highest and that on the model is lowest.

(8) The attack of the model and of the white imperfect mimics are equal, and the attack rate of the control and the other imperfect mimic are equal, and different from that of the control: $P_c = P_{ow} \neq P_o = P_w$ for group 1 and $P_c = P_o \neq P_{ow} = P_w$ for groups 2 and 3. This scenario indicates full generalisation to the white imperfect mimic but not to the other imperfect mimic if $P_c > P_w$.

(9) The attack rate of the control and the imperfect mimic with orange are equal, and the attack rates of the model and the white mimic are different and different from that of the control: $P_c = P_{ow} \neq P_o \neq P_w$ for group 1 $P_c = P_o \neq P_{ow} \neq P_w$ for groups 2 and 3. This scenario indicates partial generalisation to the white imperfect mimic, if the attack rate on the control is highest and that on the model is lowest.

**Results of the memorability and generalisation tests for the three groups.**

| **Group 1**<br>**orange as the escaping model** | | | | scenario 1<br>$P_c = P_{ow} = P_o = P_w$<br>no memory | scenario 2<br>$P_c \neq P_{ow} \neq P_o \neq P_w$<br>partial generalisation to both mimics | scenario 3<br>$P_c \neq P_{ow} = P_o = P_w$<br>full generalisation to both mimics | **scenario 4**<br>$\mathbf{P_o \neq P_c = P_{ow} = P_w}$<br>**no generalisation** | <u>**scenario 5**</u><br><u>$\mathbf{P_c = P_w \neq P_o = P_{ow}}$</u><br><u>**full generalisation same colour (orange)**</u> | scenario 6<br>$P_c \neq P_o \neq P_{ow} = P_w$<br>partial generalisation equally to both mimics | **scenario 7**<br>$\mathbf{P_c = P_w \neq P_o \neq P_{ow}}$<br>**partial generalisation same colour (orange)** | scenario 8<br>$P_c = P_{ow} \neq P_o = P_w$<br>full generalisation other colour (white) | scenario 9<br>$P_c = P_{ow} \neq P_o \neq P_w$<br>partial generalisation other colour (white) |
|---|---|---|---|---|---|---|---|---|---|---|---|---|
| | attacked | presented | attack rates | | | | | | | | | |
| control | 10 | 23 | $P_c$ | 0.250 | 0.435 | 0.435 | **0.319** | <u>**0.391**</u> | 0.435 | **0.391** | 0.304 | 0.304 |
| **orange=model** | 1 | 23 | $\mathbf{P_o}$ | 0.250 | 0.043 | 0.188 | **0.043** | <u>**0.109**</u> | 0.043 | **0.043** | 0.196 | 0.043 |
| white | 8 | 23 | $P_w$ | 0.250 | 0.348 | 0.188 | **0.319** | <u>**0.391**</u> | 0.261 | **0.391** | 0.196 | 0.348 |
| orange/white | 4 | 23 | $P_{ow}$ | 0.250 | 0.174 | 0.188 | **0.319** | <u>**0.109**</u> | 0.261 | **0.174** | 0.304 | 0.304 |
| | | | number of parameters (attack rates) | 1 | 4 | 2 | **2** | <u>**2**</u> | 3 | **3** | 2 | 3 |
| | | | log(L) - K | -22.468 | -19.694 | -21.339 | **-20.545** | <u>**-20.239**</u> | -20.091 | **-19.773** | -22.151 | -20.516 |
| | | | AICc | 47.127 | 49.610 | 47.279 | **45.690** | <u>**45.079**</u> | 47.446 | **46.809** | 48.903 | 48.296 |

| **Group 2**<br>**orange/white as the escaping model** | | | | scenario 1<br>$P_c = P_{ow} = P_o = P_w$<br>no memory | scenario 2<br>$P_c \neq P_{ow} \neq P_o \neq P_w$<br>partial generalisation to both mimics | <u>**scenario 3**</u><br><u>$\mathbf{P_c \neq P_{ow} = P_o = P_w}$</u><br><u>**full generalisation to both mimics**</u> | scenario 4<br>$P_{ow} \neq P_c = P_o = P_w$<br>no generalisation | **scenario 5**<br>$\mathbf{P_c = P_w \neq P_o = P_{ow}}$<br>**full generalisation orange mimic** | **scenario 6**<br>$\mathbf{P_c \neq P_{ow} \neq P_o = P_w}$<br>**partial generalisation equally to both mimics** | scenario 7<br>$P_c = P_w \neq P_o \neq P_{ow}$<br>partial generalisation orange mimic | scenario 8<br>$P_c = P_o \neq P_{ow} = P_w$<br>full generalisation white mimic | scenario 9<br>$P_c = P_o \neq P_{ow} \neq P_w$<br>partial generalisation white mimic |
|---|---|---|---|---|---|---|---|---|---|---|---|---|
| | attacked | presented | attack rates | | | | | | | | | |
| control | 14 | 25 | $P_c$ | 0.250 | 0.560 | <u>**0.435**</u> | 0.319 | **0.391** | **0.560** | 0.391 | 0.304 | 0.304 |
| **orange/white=model** | 2 | 25 | $\mathbf{P_{ow}}$ | 0.250 | 0.080 | <u>**0.188**</u> | 0.080 | **0.109** | **0.080** | 0.080 | 0.196 | 0.080 |
| white | 5 | 25 | $P_w$ | 0.250 | 0.200 | <u>**0.188**</u> | 0.319 | **0.391** | **0.261** | 0.391 | 0.196 | 0.200 |
| orange | 4 | 25 | $P_o$ | 0.250 | 0.160 | <u>**0.188**</u> | 0.319 | **0.109** | **0.261** | 0.160 | 0.304 | 0.304 |
| | 25 | 100 | | | | | | | | | | |
| | | | number of parameters (attack rates) | 1 | 4 | <u>**2**</u> | 2 | **2** | **3** | 3 | 2 | 3 |
| | | | log(L) - K | -24.422 | -20.681 | <u>**-21.566**</u> | -23.116 | **-22.407** | **-21.109** | -22.226 | -23.368 | -22.803 |
| | | | AICc | 51.034 | 51.584 | <u>**47.732**</u> | 50.831 | **49.415** | **49.481** | 51.715 | 51.337 | 52.868 |

| **Group 3**<br>**orange/white as the unpalatable model** | | | | scenario 1<br>$P_c = P_{ow} = P_o = P_w$<br>no memory | scenario 2<br>$P_c \neq P_{ow} \neq P_o \neq P_w$<br>partial generalisation to both mimics | scenario 3<br>$P_c \neq P_{ow} = P_o = P_w$<br>full generalisation to both mimics | scenario 4<br>$P_{ow} \neq P_c = P_o = P_w$<br>no generalisation | <u>**scenario 5**</u><br><u>$\mathbf{P_c = P_w \neq P_o = P_{ow}}$</u><br><u>**full generalisation orange mimic**</u> | scenario 6<br>$P_c \neq P_{ow} \neq P_o = P_w$<br>partial generalisation equally to both mimics | scenario 7<br>$P_c = P_w \neq P_o \neq P_{ow}$<br>partial generalisation orange mimic | scenario 8<br>$P_c = P_o \neq P_{ow} = P_w$<br>full generalisation white mimic | scenario 9<br>$P_c = P_o \neq P_{ow} \neq P_w$<br>partial generalisation white mimic | scenario10<br>$P_{ow} = P_o \neq P_c \neq P_w$<br>full generalisation orange white mimic most attacked |
|---|---|---|---|---|---|---|---|---|---|---|---|---|---|
| | attacked | presented | attack rates | | | | | | | | | | |
| control | 6 | 18 | $P_c$ | 0.250 | 0.333 | 0.435 | 0.319 | <u>**0.391**</u> | 0.333 | 0.391 | 0.304 | 0.304 | 0.333 |
| **orange/white=model** | 1 | 18 | $\mathbf{P_{ow}}$ | 0.250 | 0.056 | 0.188 | 0.056 | <u>**0.109**</u> | 0.056 | 0.056 | 0.196 | 0.056 | 0.109 |
| white | 10 | 18 | $P_w$ | 0.250 | 0.556 | 0.188 | 0.319 | <u>**0.391**</u> | 0.261 | 0.391 | 0.196 | 0.556 | 0.556 |
| orange | 1 | 18 | $P_o$ | 0.250 | 0.056 | 0.188 | 0.319 | <u>**0.109**</u> | 0.261 | 0.056 | 0.304 | 0.304 | 0.109 |
| | 18 | 72 | | | | | | | | | | | |
| | | | number of parameters (attack rates) | 1 | 4 | 2 | 2 | <u>**2**</u> | 3 | 3 | 2 | 3 | 3 |
| | | | log(L) - K | -17.584 | -13.701 | -17.650 | -16.286 | <u>**-14.458**</u> | -16.354 | -14.186 | -18.345 | -15.234 | -13.973 |
| | | | AICc | 37.358 | 37.623 | 39.901 | 37.173 | <u>**33.517**</u> | 39.972 | 35.636 | 41.289 | 37.732 | 35.209 |

Scenario numbering correspond to that mentioned above. For each scenario, the hypothesis tested is indicated and an interpretation is given.
The table presents the number of attacked and presented prey of each type, and estimates of attack rates under each scenario are given. Ln-likelihood and AICc are given for each scenario
The best scenario is indicated by bold and underlined formatting, and the alternative scenarios are indicated in bold. Note that for group 3 an additional scenario was tested, where the attack rate of the white mimic was allowed to differ from the others.
This was to account for the higher observed attack rate on the white prey.
